# Randomly primed, strand-switching MinION-based sequencing for the detection and characterization of cultured RNA viruses

**DOI:** 10.1101/2019.12.16.875872

**Authors:** Kelsey T. Young, Kevin K. Lahmers, Holly S. Sellers, David E. Stallknecht, Rebecca L. Poulson, Jerry T. Saliki, S. Mark Tompkins, Ian Padykula, Chris Siepker, Elizabeth W. Howerth, Michelle Todd, James B. Stanton

## Abstract

RNA viruses rapidly mutate, which can result in increased virulence, increased escape from vaccine protection, and false negative detection results. Targeted detection methods have a limited ability to detect unknown viruses and often provide insufficient data to detect coinfections or identify antigenic variants. Random, deep sequencing is a method that can more fully detect and characterize RNA viruses and is often coupled with molecular techniques or culture methods for viral enrichment. Viral culture coupled with third-generation sequencing were tested for the ability to detect and characterize RNA viruses. Cultures of bovine viral diarrhea virus, canine distemper virus, epizootic hemorrhagic disease virus, infectious bronchitis virus, two influenza A viruses, and porcine respiratory and reproductive syndrome virus were sequenced on the MinION platform using a random, reverse primer in a strand-switching reaction, coupled with PCR-based barcoding. Reads were taxonomically classified and used for reference-based sequence building using a stock personal computer. This method accurately detected and identified complete coding sequence genomes with a minimum of 20× coverage depth for all seven viruses, including a sample containing two viruses. Each lineage-typing region had at least 26× coverage depth for all viruses. Furthermore, analyzing the canine distemper virus sample through a pipeline devoid of canine distemper virus reference sequences modeled the ability of this protocol to detect unknown viruses. These results show the ability of this technique to detect and characterize dsRNA, negative- and positive-sense ssRNA, nonsegmented, and segmented RNA viruses.

## Introduction

RNA viruses are common etiological agents of animal diseases. Many of these diseases, such as influenza,^1^ Newcastle disease (ND),^2^ epizootic hemorrhagic disease (EHD),^3^ infectious bronchitis (IB),^4^ and porcine respiratory and reproductive syndrome (PRRS)^5^ are global economic and/or health burdens for domestic and wild animal populations. Moreover, RNA viruses account for most emerging diseases due to the swift production of genetic variants, enabling rapid evolution for adapting to environments and hosts.^6^ The low fidelity of the viral-encoded RNA-dependent RNA polymerases significantly contributes to the genetic diversity, causing mutations up to a million times higher compared to cells.^7^ Additionally, genetically related viral species with segmented genomes (e.g., reoviruses and orthomyxoviruses) can reassort genomic segments, resulting in increased genetic variation. Enhanced virulence, resistance to vaccines, and ability for novel tissue tropism can also occur due to nucleotide mutations, and reassortment events.^8-10^ Therefore, there is a need for rapid detection of variants through whole genome sequencing for proper diagnosis, treatment, control, and prevention.^11^

The molecular detection of RNA viruses is traditionally done using methods such as polymerase chain reaction (PCR), real-time PCR (rtPCR), cloning, and *in situ* hybridization (ISH).^12,13^ However, these methods use targeted approaches that require prior knowledge of the viral genome for detection and are inefficient for the discovery of novel viruses, mixed infections, and identifying whole genomes. The advent of next generation sequencing (NGS) has permitted new techniques that circumvent some issues with targeted RNA sequencing. Platforms like Illumina allow untargeted deep sequencing for novel virus detection; but can be expensive and labor intensive, particularly for whole genome sequencing.^14^ Major drawbacks of these platforms is the generation of large volumes of raw data and short reads which require high performance computers and extensive computational analysis.^15^ Another limitation of untargeted RNA sequencing is the low relative abundance of viral RNA compared to host cellular RNA, which may require depletion of ribosomal RNA (rRNA) and/or enrichment of viral RNA to obtain an ample number of reads for viral strain detection.^16,17^ For these reasons, using NGS for accurate, whole viral genome sequencing remains challenging.

The long read sequencing technology provided by MinION sequencing (Oxford Nanopore Technologies (ONT), Oxford, UK) has enabled rapid, inexpensive, high-throughput, whole-genome sequencing of viruses.^18^ MinION-based viral metagenomics studies have accurately sequenced and identified whole genomes of chikungunya virus, Zaire ebolavirus, hepatitis C virus, Venezuelan equine encephalitis virus, and Zika virus using untargeted approaches by targeting poly-A sites and directly sequencing RNA (dRNA-seq) or using primer-extension pre-amplification method (Round A/B); however, both of these methods resulted in low quality reads and poor depth of coverage across the viral genome.^19,20^ Other methods such as sequence independent primer amplification (SISPA) have been used with MinION to obtain whole genome sequences for bovine enterovirus from culture and canine distemper virus from the brain of an affected dog.^21,22^ While virus targeting can be accomplished with the previously mentioned rRNA depletion or sequence targeting, it is also possible to couple this newest sequencing technology with classical virus culture for viral enrichment. The aim of this study was to develop a method to simultaneously detect and characterize various RNA viruses from culture by using a randomly primed, strand-switching approach and sequencing on MinION. Viruses were selected to include a double-stranded RNA virus (epizootic hemorrhagic disease virus 2 [EHDV-2; family *Reoviridae*, genus *Orbivirus*]), positive-stranded viruses (infectious bronchitis virus [IBV; family *Coronaviridae*, genus *Gammacoronavirus*, species *Avian coronavirus*], porcine reproductive and respiratory syndrome virus [PRRSV; family *Arteriviridae*, genus *Betaarterivirus*, species *Betaartervirus suid 2*], and bovine viral diarrhea virus [BVDV; family *Flaviviridae*, genus *Pestivirus*, species *Pestivirus B*]), and negative-stranded viruses (canine distemper virus [CDV; family *Paramyxoviridae*, genus *Morbillivirus*, species *Canine morbillivirus*] and influenza A viruses [IAV; family *Orthomyxoviridae*, genus *Alphainfluenzavirus*, species *Influenza A virus*] isolated from a dog and from a pig). Segmented (EHDV-2 and IAV) and unsegmented genomes (BVDV, CDV, and PRRSV) were also represented in this study. This approach provides rapid, complete genome coding sequences (CDS) for unknown RNA viruses in culture fluids and demonstrates the utility of the random hexamer-based strand-switching primer for MinION library synthesis.

## Materials and methods

### Samples

The EHDV-2 isolate was propagated on cattle pulmonary artery endothelial cells (CPAE) from spleen/lung tissue from a white-tailed deer (*Odocoileus virginianus*) collected in Georgia, USA in 2016 at the Southeastern Cooperative Wildlife Disease Study at the University of Georgia (UGA); CPAE cells are also persistently infected with BVDV. The CDV sample was isolated from the brain of an infant, female raccoon (*Procyon lotor*) from Kentucky in 2018 using African green monkey kidney cells expressing canine signaling lymphocytic activation molecule (Vero-Dog SLAM cell line) in the Athens Veterinary Diagnostic Laboratory (AVDL). The canine-origin IAV sample was collected from a nasal swab of a 7-year-old, male, boxer dog in 2015 from Georgia and was cultured in embryonated chicken eggs at the Poultry Diagnostic and Research Center (PDRC), UGA. The Center for Vaccines and Immunology, UGA provided the swine-origin influenza sample, which was isolated in 2019 from a 6-month-old, female, Hampshire-cross pig from Georgia after testing positive for IAV by immunohistochemistry and PCR. Swine IAV from lung homogenate was propagated on Madin-Darby canine kidney (MDCK) cells. The IBV sample (Mass vaccine) was cultured in embryonated chicken eggs and supplied by the PDRC. Isolate VR2385 of PRRSV was cultured on MARC-145 cells at the Veterinary Diagnostic Laboratory, Iowa State University. Reverse-transcription real-time PCR (RT-rtPCR) was conducted for BVDV, EHDV-2,^3^ CDV, canine-origin influenza, and IBV (Table 1). The CDV (AVDL), canine IAV (PDRC), and IBV (PDRC) RT-rtPCR assays were done using in-house diagnostic methods. For BVDV, (RT-rtPCR) was performed using a modified protocol^23^ with SuperScript® III First-Strand Synthesis SuperMix for qRT-PCR (Invitrogen, Carlsbad, CA) and amplified in a CFX96™ Touch Real-Time PCR Detection System (Bio-Rad Laboratories, Inc, Hercules, CA) thermal cycler.

**Table 1.** Sample background information and reverse-transcription real-time PCR (RT-rtPCR) results.

| Virus | Original Host | Propagation System | RT-rtPCR<br>(Ct) | Collection |  |
| --- | --- | --- | --- | --- | --- |
|  |  |  |  | State (USA) | Year |
| BVDV | BVDV positive CPAE cells | CPAE | 24.90 | N/A | N/A |
| CDV | North American raccoon | Vero-Dog SLAM | 20.35 | Kentucky | 2018 |
| EHDV | White-tailed deer | CPAE | 15.15 | Georgia | 2016 |
| IBV | N/A | ECE | 12.48 | N/A | N/A |
| Canine-origin IAV | Dog | ECE | 16.94 | Georgia | 2015 |
| Swine-origin IAV | Hampshire-cross pig | MDCK | ND | Georgia | 2019 |
| PRRSV | N/A | MARC-145 | ND | N/A | N/A |

BVDV= bovine viral diarrhea virus; CDV = canine distemper virus; Ct = cycle threshold; CPAE = cattle pulmonary artery endothelial; ECE = embryonated chicken eggs; EHDV = epizootic hemorrhagic disease virus; IAV = influenza A virus; IBV = infectious bronchitis virus; MDCK = Madin-Darby Canine Kidney; N/A = not applicable; ND = not determined; PRRSV = porcine reproductive and respiratory syndrome virus.

### Total RNA extraction

All RNA extraction to sequencing steps for all viruses, except for PRRSV and a CDV replicate, were conducted at UGA. Library preparation and sequencing of PRRSV and a CDV replicate were conducted at Virginia Tech University. Total RNA for CDV, EHDV-2, BVDV, swine IAV, and canine IAV was extracted using 1 ml of culture supernatant using Trizol® LS Reagent (ThermoFisher Scientific, Waltham, MA) following manufacturer’s protocol. Total RNA was eluted in 88.5 µl of nuclease-free water (Qiagen, Hilden, Germany). DNase treatment was performed using RNase-Free DNase Set (Qiagen) and then purified using RNeasy® MinElute® Cleanup Kit (Qiagen) per manufacturer’s instructions. Total RNA for IBV was extracted using the QIAamp® Viral RNA Mini Kit (Qiagen) following manufacturer’s protocol. At Virginia Tech, RNA was extracted from PRRSV and one replicate of CDV using the QIAamp Viral RNA Mini Kit (Qiagen) according to the manufacturer’s instructions. Concentrations were measured using the Qubit® RNA HS Assay Kit (ThermoFisher Scientific) on a Qubit® 3.0 Fluorometer.

### Strand-switching cDNA synthesis

Strand-Switching cDNA synthesis for MinION sequencing was completed by modifying the 1D PCR barcoding cDNA (SQK-LSK108) protocol from ONT. Reverse transcription was performed by combining 8 µl of total RNA, 2 µl of 1 µM PCR-RH-RT primer (5’ - /5Phos/<u>ACTTGCCTGTCGCTCTATCTTC</u>NNNNNN-3’; synthesized by Integrated DNA Technologies [IDT], Coralville, IA, with standard desalting, ONT adapter sequence is underlined), and 1 µl of 10 mM dNTPs. The reaction mixture was incubated at 65°C for 5 minutes, then snap cooled in an ice-water slurry for 1 minute. Then 4 µl of 5× RT buffer, 1 µl of 100mM DTT, 1 µl of 40 U/µl RNaseOUT™ (Invitrogen), and 2 µl of 10 µM strand-switching oligo (PCR_Sw_mod_3G) (5’ - TTTCTGTTGGTGCTGATATTGCTGCCATTACGGCCmGmGmG-3’; ONT provided sequence, synthesized by IDT with HPLC purification) were added and incubated at 42°C for 2 minutes. SuperScript™ IV Reverse Transcriptase (Invitrogen) was added at a volume of 1 µl and the reaction was incubated at the following conditions: 30 minutes at 50°C, 10 minutes at 42°C, and 10 minutes at 80°C. The cDNA was purified by KAPA Pure Beads (Kapa Biosystems, Wilmington, MA) or with AMPure XP beads (Beckman Coulter, Indianapolis, IN) at 0.7× beads: solution ratio.

### Barcoding PCR

The reverse-transcribed cDNA was amplified following ONT’s 1D PCR barcoding cDNA (SQK-LSK108) protocol using the PCR Barcoding Expansion 1-12 (EXP-PBC001) kit (ONT) and LongAmp Taq 2× Master Mix (New England Biolabs, MA) with the following thermocycling conditions: 95°C for 3 minutes; 18 cycles of 95°C for 15 seconds, 62°C for 15 seconds, 65°C for 22 minutes; 65°C for 23 minutes. The barcoded, amplified DNA was bead purified at a 0.8× ratio.

### MinION library preparation and sequencing

After the barcoding PCR, two to three samples were pooled by equal volume to make a total volume of 47 µl. MinION libraries were prepared from the pooled barcoded amplicons using the Ligation Sequencing Kit (SQK-LSK109) (ONT) following the 1D amplicon/cDNA by Ligation per ONT’s instructions. Briefly, the pooled samples were end prepped with NEBNext FFPE Repair Mix (New England Biolabs) and NEBNext End repair/dA-tailing Module (New England Biolabs) and bead purified at 1.0× bead volume. Then, sequencing adapters (ONT) were ligated onto the end-prepped library using NEBNext Quick T4 DNA Ligase (New England Biolabs) and bead purified (0.4× beads ratio) with the Long Fragment Buffer (LFB) (ONT). The final libraries were combined with Sequencing Buffer (SQB) and Loading Beads (LB) per ONT’s instructions and were sequenced on used or new FLO-MIN106 R9.4 flow cells (ONT), with the exception of the PRRSV and replicated CDV library, which were sequenced on a FLO-MIN107 R9.5, with the MinION Mk1b sequencer. Per the pre-run quality control check for previously used flow cells, a minimum of 1,000 active pores were required for sequencing. To maximize data generation from single flow cells, used flow cells were nuclease flushed, before or after sequencing, by adding a mixture of 10 µl DNase I and 190 µl DNase I Reaction Buffer (New England Biolabs) to the priming port and incubated for 30 minutes at room temperature. Sequencing was initiated by MinKNOW v.18.12.4–19.05.0 (ONT) using the 48-hour sequencing protocol with the live basecalling option turned off. During the run, available FAST5 files were processed as below to estimate number and length of viral reads to estimate sequencing time required to obtain sufficient genome coverage.

### Pre-processing of raw MinION sequence data

The FAST5 files produced from sequencing for IBV, CDV, swine-origin and canine-origin influenza, and EHDV-2 and BVDV libraries were processed using an in-house script that sequentially basecalled, demultiplexed, and trimmed adapters on a Macbook Pro (3.1 GHz Intel Core i7, 8GB) using OS X El Capitan v.10.11.6. In summary, reads were basecalled using the CPU version of Guppy v.2.1.3 (ONT) by defining the appropriate configurations for the flowcell (FLO-MIN106) and kit (SQK-LSK109) used for sequencing and library preparation. Calibration strand detection and filtering was enabled with the --calib_detect parameter. The basecalled fastq files with a qscore ≥7 were sorted into a “pass” folder using the --qscore-filtering parameter and used for further analysis. Then, reads were demultiplexed using Porechop v.0.2.4 (https://github.com/rrwick/Porechop) based on barcodes (-b output_dir) with -- require_two_barcodes setting enabled, 1,000,000 reads were aligned to all known adapter sets (--check_reads 1000000), adapters with 99% identity were trimmed from read ends (--adapter_threshold 99), and chimeric reads with middle adapters were removed.

The raw FAST5 files produced for PRRSV and the CDV replicate from Virginia Tech were basecalled using the GPU version of Guppy v.3.1.5 on a Dell Precision 3060 Tower (Intel Core i7, 31.3GB, NVIDIA GeForce GTX 1080) using Ubuntu 16.04 LTS. Guppy was initiated using the same parameters as above with the flowcell configuration defined as “FLO-MIN107”. The basecalled reads were then trimmed and demultiplexed using Porechop with the same parameters described previously.

### Virus classification and lineage typing

After basecalling, demultiplexing, and adapter trimming, reads were classified with Centrifuge v.1.0.4^24^ by the lowest taxonomical rank with default parameters with the addition of allowing 50 assignments (or hits) for each read (-k 50) using a custom index built for each library based on the propagation system. Custom indices were constructed using an exhaustive search for complete genome sequences of all possible viruses infecting vertebrates downloaded from NCBI’s nucleotide database, as of 15 March 2019, that included 80,550 viral reference genomes (Supplementary Table 1). Low-complexity regions in sequences were masked using dustmasker (NCBI C++ Toolkit, https://ncbi.github.io/cxx-toolkit). Dustmasked whole genomes of the species of cell line (see below), and/or the bovine genome (GCF_002263795.1_ARS-UCD1.2), to account for fetal bovine serum used in cell culture, were obtained using `centrifuge-download` and concatenated with the dustmasked vertebrate virus sequences to classify reads from host(s) cellular RNA. Using the concatenated files, indices were built by `centrifuge-build` with default parameters. The indices were then used to cluster reads based on short alignments to respective host and viral sequences to determine the presence and abundance of viral species in each sample using the standard out file. Reads to unexpected viral species were clustered and compared to the Genbank database using web-based nucleotide Basic Local Alignment Tool (BLASTn) (https://blast.ncbi.nlm.nih.gov) with default settings. Percentage of viral reads and host reads were determined by removing duplicate reads and then dividing the number of reads clustered by the number of total reads after demultiplexing.

Once the viral species was identified, custom lineage-typing Centrifuge indices were also built to further classify beyond viral species and assess for the possibility of mixed infections of the same virus species, as previously described for IBV.^25^ Briefly, for each main lineage per virus, one complete sequence of the lineage-typing region was selected and then combined with the respective genome of the cell line. Indices were constructed using the following: 20 N-terminal protease fragment (*N*^*pro*^) sequences for BVDV^26^ with the bovine genome (GCF_002263795.1_ARS-UCD1.2), 13 hemagglutinin (*H*) gene sequences for CDV^27^ with the African green monkey genome (GCF_000409795_Chlorocebus_sabeus_1.1), nine *VP2* sequences for EHDV^28^ with the bovine genome (GCF_002263795.1_ARS-UCD1.2), 32 spike 1 (*S1*) sequences for IBV^29^ with the chicken genome (GCF_000002315.4_Gallus_gallus-5.0), 18 hemagglutinin (*HA*) sequences and 11 neuraminidase (*NA*) sequences for IAV^30^ with the canine genome (GCF_000002285.3_CanFam3.1) or chicken genome (GCF_000002315.6_GRCg6a), and 18 *ORF5* sequences for PRRSV^31^ with the swine genome (GCF_000003025.6_Sscrofa11.1) (Supplementary Table 2). All sequences were dustmasked, assigned a unique taxonomy identification number, with the exception of the IAV sequences, and indices were built with default settings using `centrifuge-build`. Viral lineage typing was evaluated by aligning the individual trimmed and barcoded FASTQ files produced from Porechop to the respective custom lineage-typing indexes with Centrifuge, and reads were clustered by taxonomy identifications that represent potential lineage types in the sample. Each cluster of reads, representing potentially different lineages, were then used for reference based consensus building using Geneious v.11.1.3 (Biomatters, Aukland, New Zealand) to build a consensus for each lineage using the “Map to Reference” tool with medium sensitivity, re-iterating up to five times, and using the same reference sequences chosen for generating the custom Centrifuge lineage-typing databases. Consensus sequences derived for each potential lineage were analyzed using BLASTn with default settings. Results from BLASTn were sorted by query coverage and subject coverage. Subjects with the highest query, subject coverages, and identical bit-scores were considered the “top hit(s)” to identify the virus and lineage in each cluster. A coinfection was defined as different clusters of reads creating consensus sequences that matched to different lineages and reads could be parsed accordingly for final consensus building. If no coinfection was detected, all viral reads were used for consensus building.

**Table 2.**
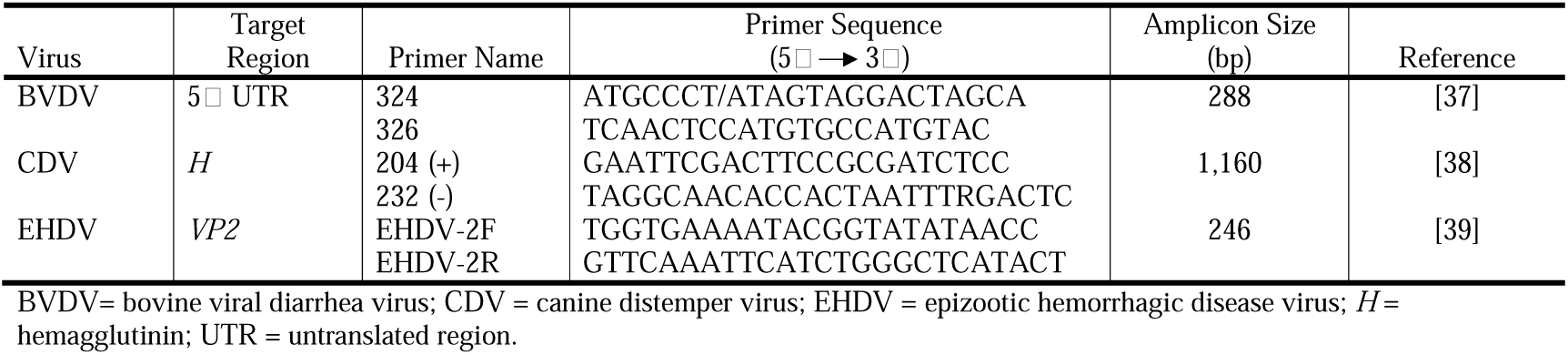
Reverse-transcription PCR (RT-PCR) primer information for Sanger sequencing.

### Viral genome consensus sequence generation

After viruses were lineage typed for each sample, all reads that aligned to respective viral species from Centrifuge output were imported into Geneious for whole-genome, reference-based consensus building with the “Map to Reference” tool, after removing duplicate reads, using the same methods from above. Consensus sequence building was done with the requirement of at least 20× coverage at each base. References used for mapping were selected from using the “top hit” result in BLASTn from the lineage-typing analysis. Due to the known errors within long homopolymer regions during MinION sequencing,^32^ consensus sequences were manually inspected and homopolymer areas that resulted in frameshifts were manually edited. The CDS regions were extracted from each edited consensus sequence and analyzed using BLASTn. For viruses with gapped genomes (e.g., CDV, IBV, and PRRSV), complete CDS and non-coding intergenic regions were extracted and analyzed with BLASTn. The “top hit” for each consensus sequence was determined by using the same criteria from above. The CDS regions for lineage-typing regions for each virus were also extracted and analyzed with BLASTn. The best “top hit”, percent pairwise identity, mean, minimum, maximum, and standard deviation of base coverage, and percentage of genome coverage were evaluated for each viral genome and lineage-typing region consensus sequences with the statistics tool in Geneious.

### Simulating novel virus identification

In addition to the above method, sequencing data from the CDV sample were analyzed in a manner that simulated novel virus identification by constructing an all vertebrate virus, custom Centrifuge index with the omission of all canine morbillivirus sequences downloaded from NCBI as of 12 April 2019 (Supplementary Table 1). The index was assembled as described previously with African green monkey genome. Trimmed and demultiplexed reads from the CDV sample were classified with Centrifuge using parameters described above. Reads were grouped by taxonomy ID and exported into Geneious to build consensus sequences as described previously. The consensus was analyzed with BLASTn by excluding canine morbillivirus (taxid: 11232) in the search set. For comparison, Kraken-style reports were created using `centrifuge-kreport` from the Centrifuge output files from the CDV-absent pipeline and the CDV-present pipeline from above. Output files were visualized using Pavian v.1.0.^33^ To discern the phylogeny of the “novel” virus among other pathogens in the *Morbillivirus* genus, the final genome CDS and phosphoprotein (*P*) gene CDS consensus sequences were aligned with 18 complete genome CDS and *P* gene CDS acquired from GenBank using the neighbor-joining algorithm in *ClustalW*^34^ with default settings. Phylogenetic trees for complete genome CDS and *P* gene CDS were inferred using the Maximum Likelihood statistical method based on the Tamura 3-parameter substitution model^35^ with bootstrap values calculated at 1000 replicates in MEGA X.^36^

### Sanger sequencing for validation

Sanger sequencing was performed for the 5’ UTR region of BVDV, *H* gene of CDV, and *VP2* segment of EHDV-2 to confirm the identity of isolates and to compare MinION consensus sequences for the lineage-typing regions. Briefly, using 8 µl of total RNA from each sample, cDNA was synthesized using SuperScript® III First-Strand Synthesis System for RT-PCR (Invitrogen) with random hexamers (50 ng/µl) following manufacturer’s instructions. The cDNA from each sample was amplified using DreamTaq Green PCR Master Mix (2×) (ThermoFisher Scientific) following manufacturer’s protocol with 10 µM of each respective primer (Table 2) targeting partial sequences of the lineage-typing regions and 1 µl of cDNA. Thermocycling conditions for PCR amplification of the 5’ UTR region for BVDV are as follows: 95°C for 5 minutes; 34 cycles of 95°C for 30 seconds, 58°C for 30 seconds, 72°C for 30 seconds; 72°C for 5 minutes. Thermocycling conditions for PCR amplification of the *H* gene for CDV are as follows: 95°C for 5 minutes; 40 cycles of 95°C for 30 seconds, 50°C for 30 seconds, 72°C for 1.5 minutes; 72°C for 10 minutes. Thermocycling conditions for PCR amplification of the *VP2* segment for EHDV are as follows: 95°C for 3 minutes; 40 cycles of 95°C for 30 seconds, 57°C for 30 seconds, 72°C for 45 seconds; 72°C for 5 minutes. Electrophoresis with a 1.0% agarose gel was performed to confirm PCR products. Amplicons were then purified using the QIAquick PCR Purification Kit (Qiagen) following manufacturer’s protocol and eluted in 30 µl nuclease-free water (Qiagen). Final concentration and purity were measured using NanoDrop™ 2000 Spectrophotometer (ThermoFisher Scientific). The purified PCR products and 5 µM of each primer (Table 2) were submitted to GENEWIZ (South Plainfield, NJ) for bidirectional Sanger sequencing.

### Sanger sequencing analysis and pairwise identity with MinION consensus sequences

Results from Sanger sequencing were analyzed with Geneious. Using the chromatograms, low quality regions (≥Q40) from the 5□ and 3□ ends were trimmed and ambiguous bases were manually edited. Primers were trimmed before aligning forward and reverse sequences from each sample using Geneious Alignment with default settings. The consensus sequence was compared using BLASTn and selecting the best “top hit” with the same criteria described above for MinION data.

Partial lineage-typing sequences from Sanger sequencing for BVDV, CDV, and EHDV were compared with the full-length lineage-typing consensus sequences from MinION using Geneious Alignment with default settings. Pairwise identity of the alignment was calculated with Geneious.

## Results

### MinION sequencing, viral classification, and lineage typing

Libraries were prepared for seven cultured samples for seven different viruses by using random hexamer primed, strand-switching for reverse transcription, PCR-based barcoding, and pooled sequencing using MinION. Sequencing occurred for a minimum of about 2 hrs and 45 min and a maximum of about 47 hrs and 37 min to obtain 567,780–6,984,000 raw reads. Total sequencing time varied between runs based on the time of day the run was started, the number of viral reads detected early in the sequencing run, and the ability to re-use a flow cell (i.e., if the flow cell was not to be reused, then it was often sequenced to near exhaustion). After the raw reads were basecalled, demultiplexed, and trimmed, 7,523–1,173,058 reads were assigned to barcodes of interest and used for viral classification (Table 3).

**Table 3.** Sequencing data from Porechop and Centrifuge.

| Virus | Estimated<br>Total Run<br>Time<br>(HH:MM) | Porechop |  | Viral<br>Reads<br>(%) | Host Reads<br>(%) |
| --- | --- | --- | --- | --- | --- |
|  |  | Total Reads | Total Bases |  |  |
| BVDV & EHDV | 8:20 | 756,373 | 299,505,703 | 12.7 | 75.2 |
| CDV | 3:20 | 183,206 | 63,116,488 | 2.10 | 80.4 |
| CDV* | 47:37 | 75,492 | 33,681,975 | 0.97 | 45.8 |
| IBV | 7:40 | 7,523 | 11,361,873 | 63.3 | 2.50 |
| Canine IAV | 2:45 | 745,382 | 341,878,915 | 36.6 | 53.2 |
| Swine IAV | 5:20 | 1,173,058 | 450,544,038 | 31.1 | 63.4 |
| PRRSV* | 47:37 | 28,418 | 18,889,591 | 57.1 | 14.3 |

BVDV= bovine viral diarrhea virus; CDV = canine distemper virus; EHDV = epizootic hemorrhagic disease virus; IAV = influenza A virus; IBV = infectious bronchitis virus; PRRSV = porcine reproductive and respiratory syndrome virus.
\* Libraries were prepared and sequenced at Virginia Tech University.

Barcoded reads for each of the seven libraries were aligned to custom-built indices with Centrifuge that detected seven different viruses belonging to three main types of RNA viral genomes: BVDV (positive-sense ssRNA), CDV (negative-sense ssRNA), EHDV-2 (dsRNA), IBV (positive-sense ssRNA), canine-origin IAV (negative-sense ssRNA), swine-origin IAV (negative-sense ssRNA), and PRRSV (positive-sense ssRNA). Additionally, three of the viruses, EHDV-2, canine-origin influenza, and swine-origin influenza, are comprised of segmented genomes. The randomly primed strand-switching method was also able to classify a sample containing EHDV-2 and BVDV. All viruses detected were categorized with viral reads representing 0.97–63.3% of all reads sequenced for each sample with percentage of host reads ranging from 2.5 to 80.4% (Table 3). Reads to unexpected viral species were analyzed with BLASTn and were determined to be short alignments to various host, often ribosomal, sequences.

Viral sequences were parsed and further classified into lineage types using custom built lineage-typing indices for Centrifuge. Reads were clustered based on the lineage-typed alignment and consensus sequences were built using Geneious. While the Centrifuge alignments suggested the possibility of 2–19 lineages per virus isolate, only one lineage per species was detected in each isolate after consensus building of each potential lineage (Table 4).

**Table 4.**
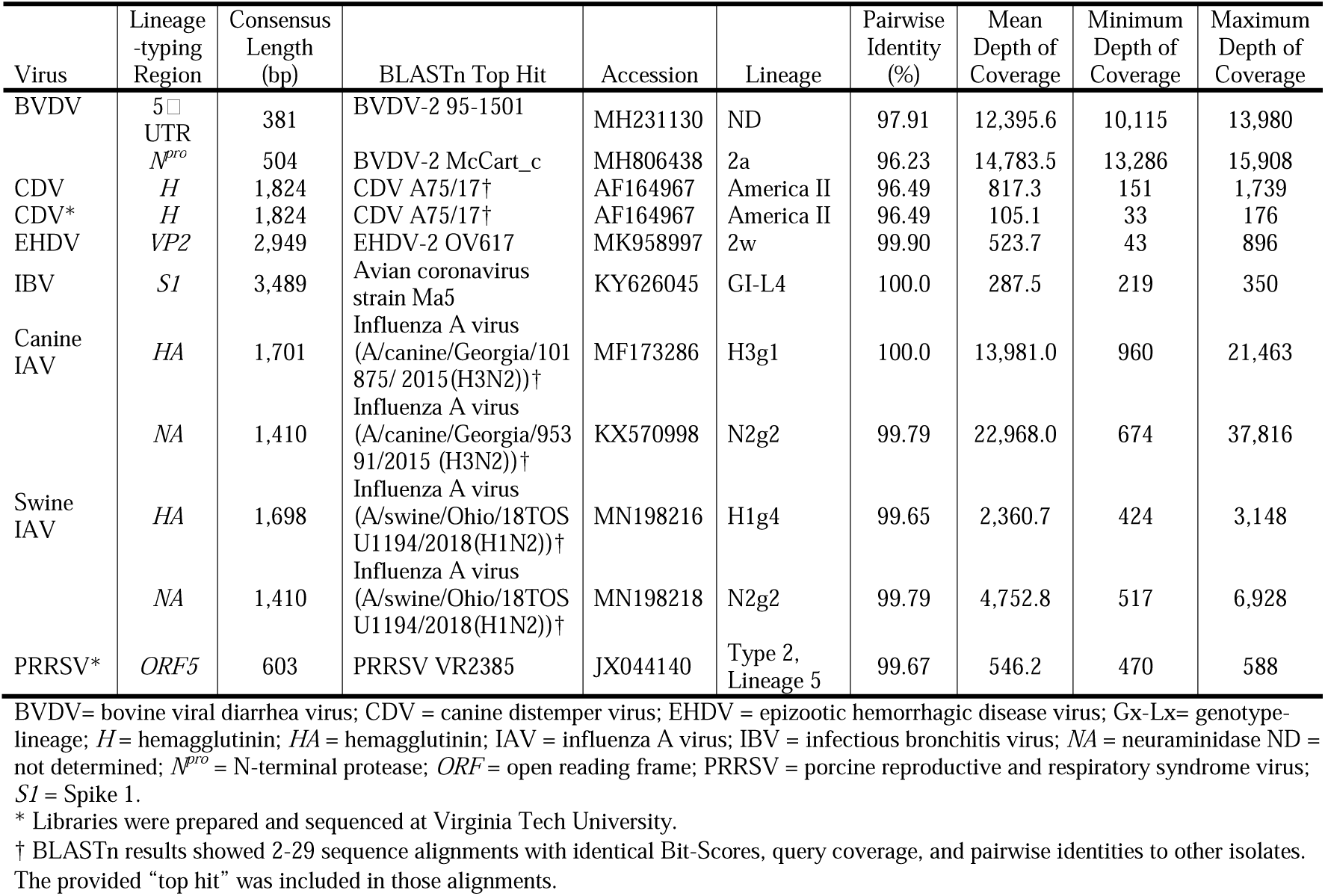
Consensus results for coding sequences (CDS) of lineage-typing regions for each virus.

The complete CDS for each virus’s lineage-typing region had at least 33× depth of coverage with a mean range of 105.1–22,968.0 (Table 4). The BLASTn comparison for the genotyping regions resulted in 96.49–100.0% pairwise identity with each “top hit” (Table 4). Using the *N*^*pro*^ region, the lineage of BVDV was determined as genotype BVDV-2a with 96.23% identity with McCart_C strain. Sequences from both libraries of CDV were genotyped as America II with 96.49% identity of the *H* gene to A75/17. The genotype for EHDV-2 was determined as EHDV-2w with 99.90% identity to the OV215 strain using the *VP2* segment. IBV was determined to be genotype 1-lineage 4, with 100.0% identity of the *S1* gene to the Ma5 strain. The *HA* and *NA* segments of the canine-origin IAV showed 100.0% and 99.79% identity, respectively, to an H3N2 subtype circulating in dogs in North America. The porcine-origin influenza was determined to be an H1N2 subtype similar to other North American porcine-origin IAVs, with 99.65% identity to the *HA* segment and 99.79% identity to the *NA* segment. The *ORF5* for PRRSV had 99.67% identity to VR2385 and was determined as a Type 2, lineage 5.

### Viral genome consensus sequence evaluation

For all seven viruses, complete genome CDS were acquired with a minimum of 26× depth of each base, with the exception of the replicated CDV with 90.2% genome coverage at 20× depth (Table 5). Additionally, complete CDS for all segments of three segmented viruses, EHDV-2 (dsRNA), swine influenza, and canine influenza (negative-sense ssRNA), were obtained with a minimum of 43× depth of coverage and an average of 99.89% identity across all segments. The mean depth for each virus was 72.1–28,014.8. The complete genome CDS consensus sequences had a high pairwise identity of 97.37–100.0% to their respective “top hit” when using BLASTn to compare sequences with GenBank databases (Table 5). The complete CDS consensus sequences for BVDV, EHDV, and canine IAV were deposited in GenBank under the following accession numbers: BVDV = MN824468; EHDV= MN824457–MN824466; canine IAV = MN812282–MN812289. The complete CDS with intergenic regions for CDV was also deposited in Genbank as accession number MN824467.

**Table 5.**
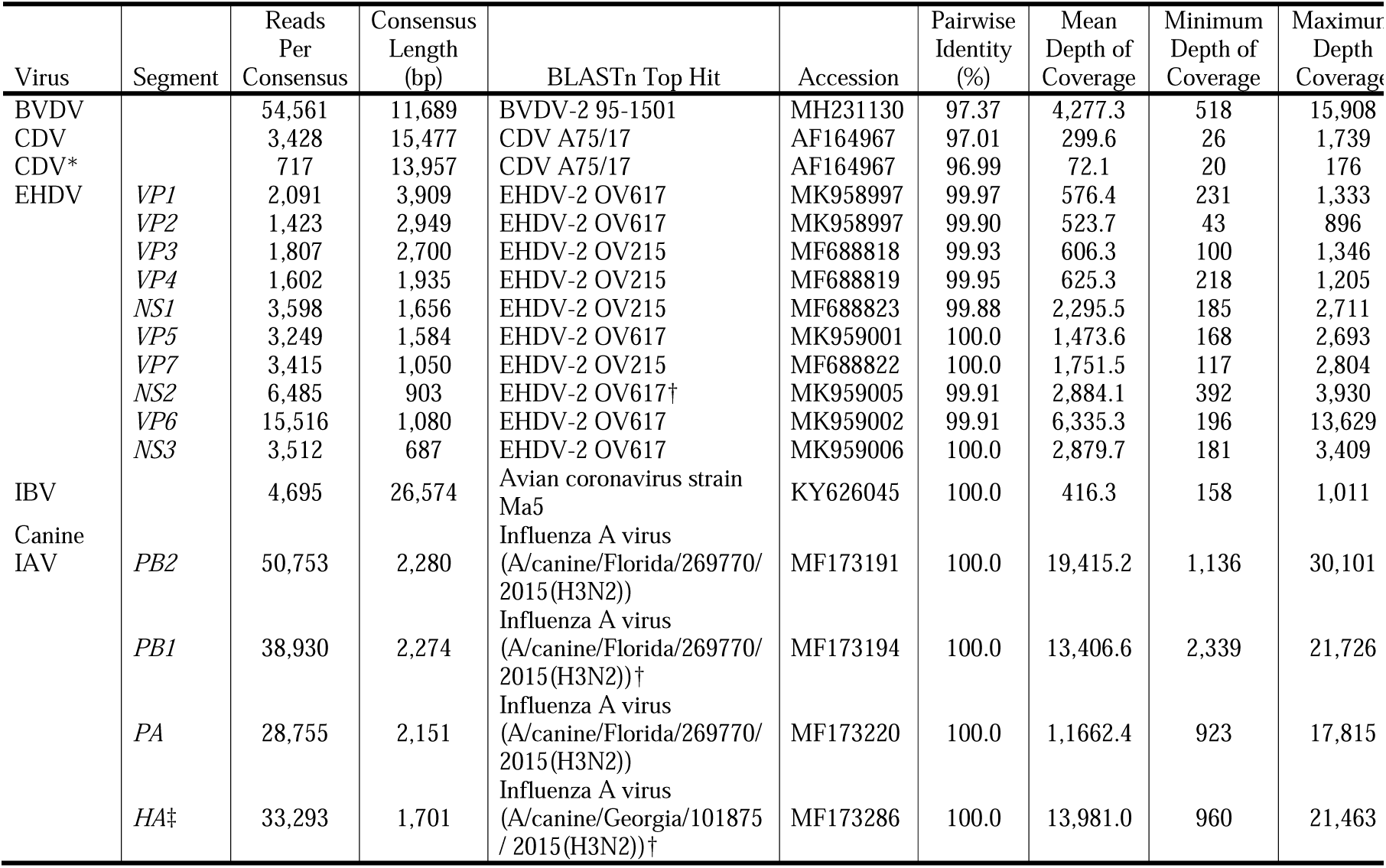
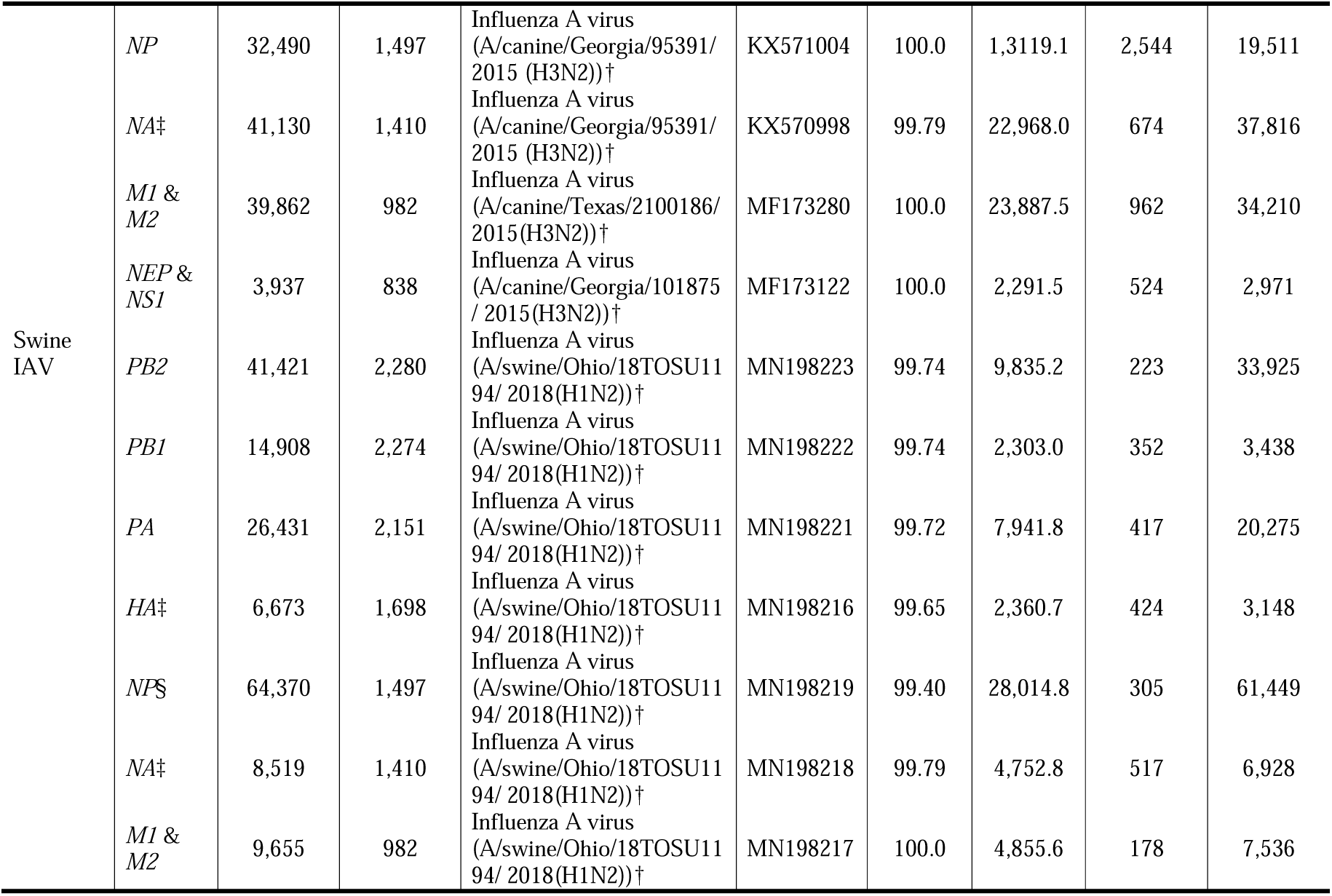
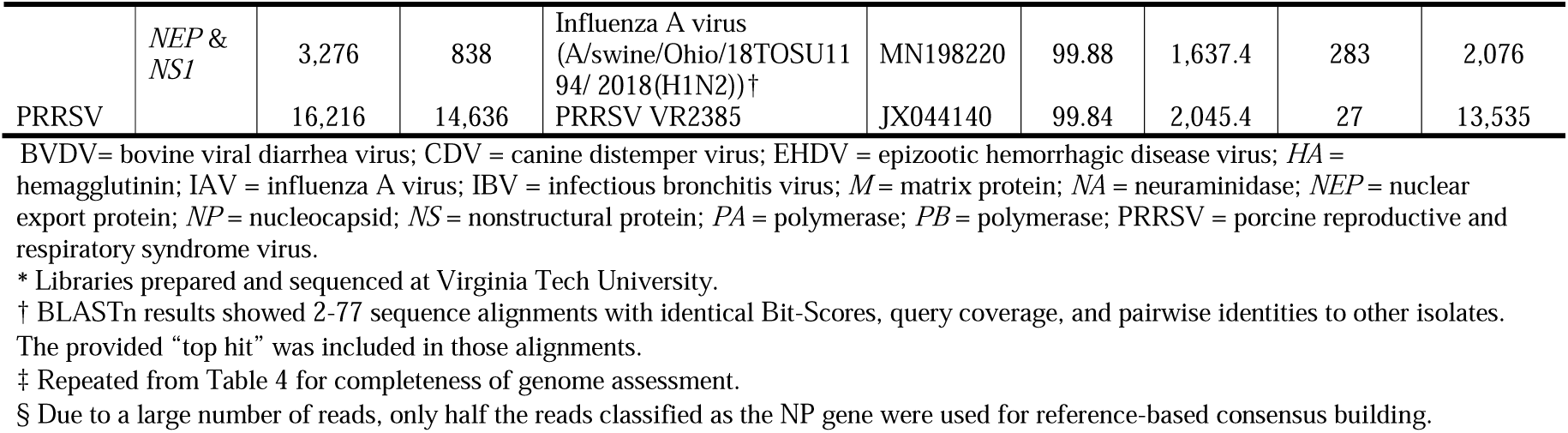
Consensus results of whole genome coding sequences (CDS) for each virus.

### Simulating novel virus identification

Under simulated conditions in which CDV would be an unknown virus, the Centrifuge standard out file for CDV analyzed showed that approximately 500 reads aligned to a morbillivirus, with 239 reads aligned to phocine distemper virus (PDV; family *Paramyxoviridae*, genus *Morbillivirus*, species *Phocine morbillivirus*) and fewer reads (<70) aligning to other more distantly related morbilliviruses: feline morbilliviruses (species *Feline morbillivirus*), rinderpest virus (species *Rinderpest morbillivirus*), peste-des-petits-ruminants virus (species *Small ruminant morbillivirus*), cetacean morbillivirus (species *Cetacean morbillivirus*), and measles virus (species *Measles morbillivirus*) (Figure 1a). In contrast, when CDV was included in the Centrifuge database, approximately 4,000 reads aligned to CDV, which was the only morbillivirus detected (Figure 1b). In the CDV-absent analysis, reads scattered across different morbilliviruses suggesting that the actual species was absent from the database but is most similar to PDV. A total of 251 (12 reads aligned to 3 or more sequences, resulting in these reads not being counted in the Centrifuge standard out file) reads were classified as PDV and were exported to Geneious for reference-based consensus building by aligning to PDV/Wadden_Sea.NLD/1988 (accession: NC_028249). The consensus sequence was analyzed with BLASTn by excluding canine morbillivirus in the search set which resulted in a 78.71% identity with PDV/Wadden_Sea.NLD/1988 (accession: KC802221.1) with 78.0% query coverage. Furthermore, phylogenetic analysis of the whole genome CDS and *P* gene CDS consensus of the dubbed “novel” sequence clustered with PDV but shows sequence divergence suggestive of a species similar to, but different from, PDV (Figure 2).

**Figure 1.**
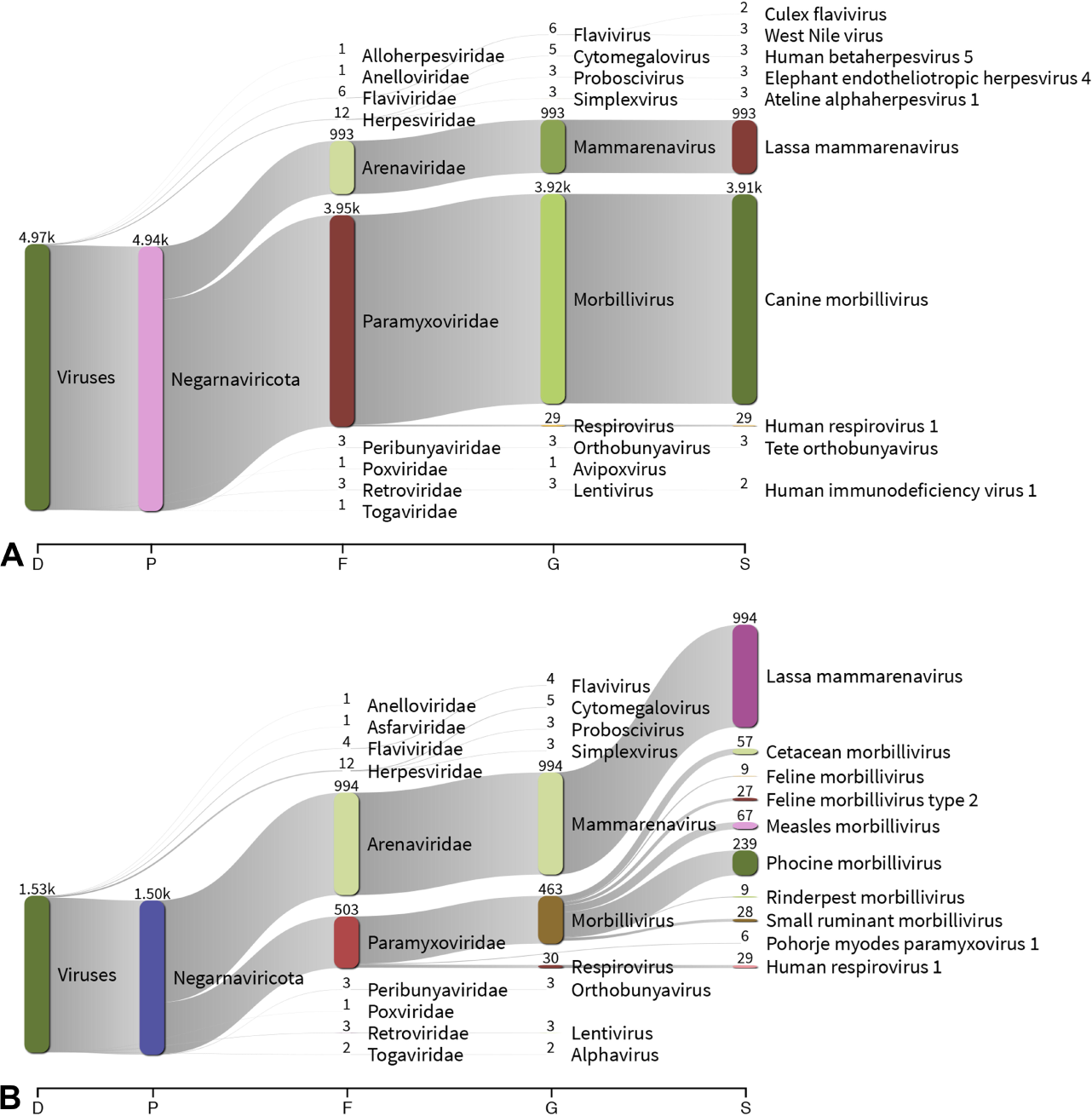
Sankey diagram displaying the top ten nodes per taxonomic classification level. Centrifuge v.1.0.4^24^ Kraken-style reports from analyzing sequences from canine distemper virus (CDV) with (a) pipeline containing CDV sequences and (b) novel virus simulation using a pipeline devoid of CDV sequences in custom Centrifuge indices were visualized with Pavian v.1.0.^33^ Number of reads assigned to taxonomic classification correspond with the width of the flow and are listed above each node. Reads hitting to the bovine and African green monkey genomes were excluded. Abbreviations are: D = domain; F = family; G = genus; P=phylum; S = species.

**Figure 2.**
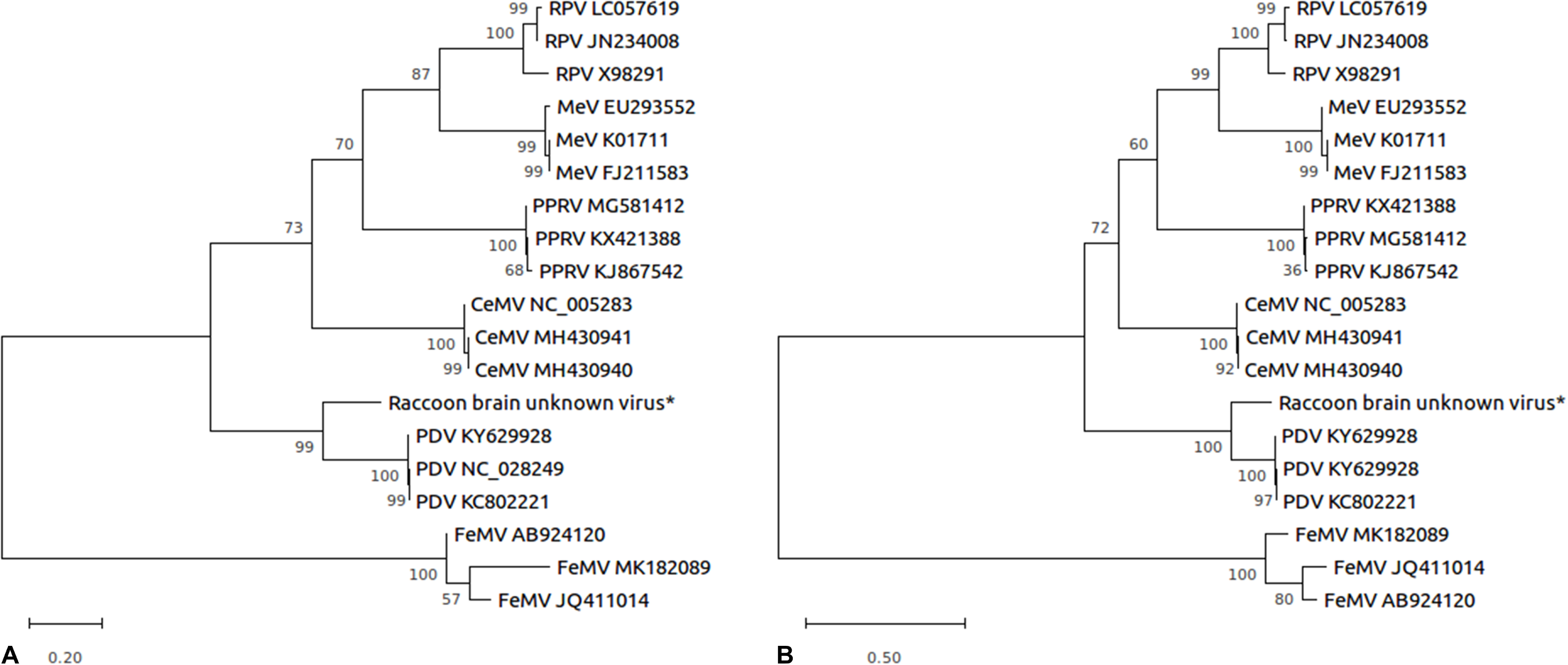
Phylogenetic analysis of morbilliviruses of (a), whole genome CDS and (b), *P* gene CDS after a data analysis of CDV sequences using a pipeline devoid of CDV reads to simulate novel virus detection and identification. Each sequence is designated by species abbreviation and GenBank accession number. The sequence denoted with an asterisk was generated from the data analysis pipeline with MinION sequencing. The evolutionary history was inferred by using the Maximum Likelihood method and Tamura 3-parameter model.^35^ These analyses involved 19 nucleotide sequences and a total of 14,578 positions (a) or 1,460 positions (b) in the final dataset. All positions containing gaps and missing data were eliminated. Evolutionary analyses were conducted in MEGA X.^36^ Abbreviations are: CeMV = cetacean morbillivirus; FeMV = feline morbillivirus; MeV = measles virus; PDV = phocine distemper virus; PPRV = peste-des-pestits-ruminants virus; RPV = rinderpest virus.

### Sanger Sequencing and pairwise identity with MinION

Sanger sequencing targeting partial sequences of the lineage-typing regions for BVDV, CDV, and EHDV was used to confirm lineage types in the samples (Table 6). For BVDV, primers targeting a partial sequence of the 5□ UTR were used and resulted in a “top hit” to BVDV-2 isolate 95-1501. Sequencing of the *H* gene for CDV resulted in a “top hit” to CDV isolate THA/VG. The *VP2* segment was targeted for EHDV and resulted in a “top hit” to EHDV-2 isolate OV617. The consensus sequences from Sanger and MinION sequencing had 100.0% pairwise identities for all three viruses (Table 6). MinION consensus sequences compared to Sanger for BVDV and EHDV had identical “top hits”. For CDV, MinION sequencing had a “top hit” to CDV A75/15 with 96.49% identity but the shorter 923 bp fragment from Sanger had a top alignment with CDV THA/VG with 96.53% identity (Table 6). The Sanger sequence is based on smaller fragments of the lineage-typing region and, therefore, the BLASTn-based pairwise identities cannot be directly compared between MinION and Sanger sequences.

**Table 6.** Sanger and MinION sequencing consensus pairwise identity results.

| Virus | Target<br>Region | MinION |  |  | Sanger |  |  | Sanger vs.<br>MinION Pairwise<br>Identity (%) |
| --- | --- | --- | --- | --- | --- | --- | --- | --- |
|  |  | Consensus<br>Length<br>(bp) | BLASTn Top Hit | Pairwise<br>Identity<br>(%) | Consensus<br>Length<br>(bp) | BLASTn Top Hit | Pairwise<br>Identity<br>(%) |  |
| BVDV | 5' UTR | 381 | BVDV-2 95-1501<br>(MH231130) | 97.91 | 268 | BVDV-2 95-1501<br>(MH231130) | 99.22 | 100.0 |
| CDV | *H* | 1,824 | CDV A75/17<br>(AF164967) | 96.49 | 923 | CDV THA/VG<br>(JX886780)* | 96.53 | 100.0 |
| EHDV | *VP2* | 2,949 | EHDV-2 OV617<br>(MK958998) | 99.90 | 158 | EHDV-2 OV617<br>(MK958998)* | 100.0 | 100.0 |

BVDV= bovine viral diarrhea virus; CDV = canine distemper virus; EHDV = epizootic hemorrhagic disease virus; *H* = hemagglutinin; UTR = untranslated region.
\* BLASTn results showed 5-12 sequence alignments with identical Bit-Scores, query coverage, and pairwise identities to other isolates. The provided “top hit” was included in those alignments.

## Discussion

A large proportion of emerging diseases are caused by RNA viruses^40^ and, due to their mutability, rapid detection is needed; however, this can be hindered due to the series of PCR panels used for identification of unknown viruses and inefficient for the discovery of coinfections.^41^ Deep sequencing-based approaches using viral nucleic acid enrichment methods have been described to address this issue, including targeted and untargeted library preparations, like SISPA.^21,22^ The methodology in this study demonstrates the application of culture-based viral enrichment followed by random, strand-switching MinION sequencing for accurately detecting and characterizing RNA viruses. RNA viruses with varying genome compositions (single stranded [positive- and negative-sense], double stranded, and segmented) were used to demonstrate the ability of untargeted strand-switching to obtain complete CDS of genotyping regions and whole genomes with viral culture enrichment methods. Moreover, two viruses (EHDV-2 and BVDV) were detected from a single sample, illustrating the utility of the random sequencing approach. Lastly, data analysis for one sample (CDV) was treated as a novel virus, highlighting the feasibility of this method to identify a new or poorly characterized virus.

This random sequencing approach proved to be robust across various RNA viruses in obtaining full length CDS of the complete genome after using an unbiased, fast aligner to identify the likely virus, followed by reference-based consensus building to identify the lineage type. With at least 26× depth of coverage across the genome for all viruses, whole genome CDS had 96.99–100.0% identity to the top BLASTn hits. Complete CDS for all segments of three segmented viruses, EHDV-2 (dsRNA), swine influenza, and canine influenza (negative-sense ssRNA), were obtained with a minimum of 43× depth of coverage and an average of 99.89% identity across all segments. The best “top hit” for each segment does not match across all segments to the same isolate, which may be due to reassortment events for segmented genomes^8,9^ and the unavailable sequence data for all segments for some isolates in NCBI.

The alignments of the whole genome CDS and the lineage-typing regions were similar and consistent with the origin of the samples in this study. The identification of the EHDV as an EHDV-2w with highest similarity to EHDVs circulating in white-tailed deer in the southeastern United States in 2017 is consistent with the collection of this sample from a white-tailed deer in Georgia in 2016. The influenza sample of canine origin was determined as a canine H3N2 strain, matching the typing completed as part of the original diagnostic case workup. The full sequencing provided by this method was able to confirm that the isolate in this study was most similar to canine H3N2 viruses circulating in the southeastern United States in 2015, consistent with the time and geographic location in which this sample was collected. Lineage typing of the swine-origin influenza virus categorized it as an H1N2 most similar to an H1N2 IAV isolated from a swine in 2018 from Ohio. It is possible that the slightly lower percent identity for the swine-origin influenza is due to a relative paucity of sequence data available for 2019 swine-origin IAV at this time. The CDV America II lineage, the classification of CDV in this study, is a common lineage found in North American wildlife^42^ and is consistent with the collection of this sample from a raccoon in Kentucky. The sequences for IBV and PRRSV were highly similar to the known sequences of those isolates.

This protocol was also tested on two scenarios. The first was the detection of two viruses in a single culture system, demonstrating its ability to simultaneously obtain accurate, complete CDS for rapid detection and characterization of viruses in coinfected samples, comparable to other studies using advanced sequencing for analysis of cultured viruses.^43^ The second was the data analysis under the simulating conditions of CDV being a novel virus. This resulted in read alignments that spanned across the *Morbillivirus* genus. Excluding the background hits, the largest proportion of reads hit to PDV and consensus building with these reads had a low pairwise identity (78.71%), suggesting it was not a PDV. The phylogenetic divergence gives evidence that the virus in the sample belongs to the *Morbillivirus* genus and is phylogenetically related to PDV. If CDV was truly an unknown virus, the Centrifuge output would have suggested the identification of a novel or divergent virus that is most similar to PDV sequences, consistent with the known close genetic relationship between CDV and PDV.^44^ Furthermore, in the event of an unknown viral etiology and due to the multiplexing capabilities of MinION sequencing, DNA library approaches can be applied and sequenced concurrently with the random strand-switching library to identify RNA or DNA virus in the sample.^45,46^

Sanger sequencing was performed on partial lineage-typing sequences for BVDV, CDV, and EHDV and compared with the full-length lineage-typing regions obtained from MinION sequencing, resulting in 100.0% pairwise identity. The 100% pairwise identity between the Sanger and MinION sequences is consistent with previous results using MinION sequencing,^2,25^ that demonstrated the ability to obtain accurate MinION sequences by increasing depth of coverage.^47,48^ Sanger sequencing and BLASTn results for the 5□ UTR for BVDV and *VP2* for EHDV matched the MinION lineage-typing region BLASTn results. For the *H* gene for CDV, Sanger sequencing had a top BLASTn alignment with isolate THA/VG with 96.53%; however, the results did show 96.0% identity with isolate A75/15, comparable to MinION sequencing result of 96.49% identity to isolate A75/15. It is noteworthy that aligning the Sanger and MinION results for the *H* gene in CDV showed 100.0% pairwise identity. The differences in BLASTn top alignments are attributed to the shorter sequence from Sanger sequencing, causing a slightly lower sequence specificity. Longer fragments, particularly whole genome sequencing, have shown improved resolution for genotyping analyses.^49,50^ Additionally, the limitations of partial sequences used in Sanger sequencing are illustrated by the inability of Sanger sequencing to identify a single best hit for EHDV-2 and CDV, which resulted in multiple hits with identical BLASTn scores; whereas, the MinION-based method obtained complete CDS of the *VP2* segment and *H* gene allowing for a single best hit for these viruses. In addition to more accurate classification, whole genome sequencing of viruses is advantageous in providing more data for detection, epidemiology, genotyping and phylogenetic analysis compared to partial and/or targeted sequences obtained from other classical sequencing methods. The relatively limited data provided by targeted sequencing may require further sequencing and expenses to obtain additional biological information. In particular, different genetic sequences are used for phylogenetic analysis and genotyping. Some viruses also have multiple genotyping regions, such as BVDV, EHDV, and influenza.^26,28,30^ Furthermore, recombination events are often difficult to identify with partial sequences and are important in investigating host range, virulence, and vaccine evasion.^51-53^ Thus, routine, whole genome sequencing of viruses gives quick turnaround data for various analyses and could provide more comprehensive databases needed to understand viral evolution.

Random, deep sequencing for many high-throughput sequencing platforms has some caveats such as false hits. As one example, illustrated in Figure 1, some reads were classified as lassa mammarenavirus, various herpesviruses, and others. Endogenous retroviruses and some DNA viruses were also frequently found to be present after annotating sequences. After clustering and aligning these reads to GenBank using BLASTn, it was determined that the reads were short sequences that matched to various host sequences. Reference sequences of the host and propagation systems of the cultured viruses were included in the Centrifuge indices to reduce these misalignments; however, these genomes are not as well described as other commonly studied organisms and may not represent the full sequence diversity of the host genome. Users can adjust alignment settings to help reduce the number of false hits, but as with any test, increasing the specificity of the test will negatively impact sensitivity. Similar to other fields (e.g., background lesions in pathology, growth of nonpathogenic bacteria in bacterial cultures), confirmatory tests may be required and analysis of deep sequencing results requires evaluation by a trained user, one experienced with bioinformatic methodology and with knowledge of veterinary infectious diseases.

While rapid, cost-effective whole-genome sequencing may be useful in a diagnostic setting, ease of use and robustness across laboratories is key for deployment. For this reason, libraries for CDV and PRRSV were prepared at different laboratories. Both libraries for CDV had similar results; however, the differences can possibly be attributed to the use of a sequencing kit intended for 1D sequencing with a FLO-MIN107 (R9.5) flowcell that is typically used for 1D^2^ sequencing. The repercussions of the 1D kit with a 1D^2^ flowcell combination are not well known but it is of interest to note that trimming the basecalled files resulted in 1,691,792 reads removed due to middle adapters, significantly decreasing the total number of reads assigned to each barcode.

The present study provides promising results for quick identification of unknown cultured RNA viruses by using MinION sequencing with a previously untested random approach. Future studies to compare the varying random methods of deep sequencing are required to determine the most efficient methods. As with other deep sequencing methods,^50^ MinION-based sequencing will also likely be used to metagenomically detect viruses directly from clinical samples (e.g., serum, swabs, tissues), and work is ongoing to investigate this usage.

MinION sequencing is a cost-effective way to multiplex samples and achieve long reads for more accurate genome consensus building compared to other short read sequencing technologies. Overall, the addition of full-genome sequencing to more routine diagnostic use will increase the available knowledge regarding sequence diversity and allow for improved tracking of viruses and a better understanding of the genetic determinants of viral pathogenesis.

## Acknowledgments

The authors acknowledge Vanessa Gauthiersloan, Samantha Day and Erich Linnemann from the Poultry Diagnostic & Research Center, University of Georgia, Clara Kienzle-Dean from the Southeastern Cooperative Wildlife Disease Study, University of Georgia, and Pablo Pinyero from the Veterinary Diagnostic Lab, Iowa State University for their technical help.

## Declaration of conflicting interests

The authors declared no potential conflicts of interest with respect to the research, authorship, and/or publication of this article.

## Funding

The project described was supported by Agriculture and Food Research Initiative Competitive Grant no. 2018-67015-28306 from the USDA National Institute of Food and Agriculture. This project has been funded in part with funds from the National Institute of Allergy and Infectious Diseases, National Institutes of Health, Department of Health and Human Services, under Center of Excellence of Influenza Research and Surveillance (CEIRS) under contract number HHSN272201400004C (SMT).

## Supplementary material

Supplementary material for this article is available online.

