## Supplementary Table for "Randomly primed, strand-switching MinION-based sequencing for the detection and characterization of cultured RNA viruses"

**Supplementary Table 1.** Search strings used to obtain all vertebrate virus sequences to build custom Centrifuge database to taxonomically classify sequencing reads.

| **Centrifuge Index** | **Number of Sequences** | **Search String** | **Date of Download** |
| --- | --- | --- | --- |
| all vertebrate virus index | 80,550 | (txid10292[ORGN] OR txid548682[ORGN] OR txid10508[ORGN] OR txid151340[ORGN] OR txid151341[ORGN] OR txid137992[ORGN] OR txid10486[ORGN] OR txid10240[ORGN] OR txid687329[ORGN] OR txid39724[ORGN] OR txid1542744[ORGN] OR txid10780[ORGN] OR txid10993[ORGN] OR txid585893[ORGN] OR txid10880[ORGN] OR txid291484[ORGN] OR txid76803[ORGN] OR txid11118[ORGN] OR txid12058[ORGN] OR txid39733[ORGN] OR txid11974[ORGN] OR txid11050[ORGN] OR txid12283[ORGN] OR txid11018[ORGN] OR txid178830[ORGN] OR txid1513294[ORGN] OR txid11270[ORGN] OR txid11266[ORGN] OR txid11158[ORGN] OR txid11617[ORGN] OR "Peribunyaviridae"[Organism] OR txid11308[ORGN] OR txid11632[ORGN] OR txid10404[ORGN]) NOT phage[All Fields] NOT patent[All Fields] NOT unverified[Title] NOT vector[Title] NOT synthetic[Title] NOT chimeric[Title] NOT miRNA NOT txid28384[ORGN] NOT method[Title] NOT "partial"[Title] NOT "region"[Title] NOT "uncultured virus"[Organism] AND 00000000101[SLEN] : 00001300000[SLEN] AND ("complete genome" OR "complete segment" OR "whole genome" OR "full genome" OR "whole segment" OR "full segment") | 15 March 2019 |
| all vertebrate virus index without CDV sequences | 84,729 | (txid10292[ORGN] OR txid548682[ORGN] OR txid10508[ORGN] OR txid151340[ORGN] OR txid151341[ORGN] OR txid137992[ORGN] OR txid10486[ORGN] OR txid10240[ORGN] OR txid687329[ORGN] OR txid39724[ORGN] OR txid1542744[ORGN] OR txid10780[ORGN] OR txid10993[ORGN] OR txid585893[ORGN] OR txid10880[ORGN] OR txid291484[ORGN] OR txid76803[ORGN] OR txid11118[ORGN] OR txid12058[ORGN] OR txid39733[ORGN] OR txid11974[ORGN] OR txid11050[ORGN] OR txid12283[ORGN] OR txid11018[ORGN] OR txid178830[ORGN] OR txid1513294[ORGN] OR txid11270[ORGN] OR txid11266[ORGN] OR txid11158[ORGN] OR txid11617[ORGN] OR "Peribunyaviridae"[Organism] OR txid11308[ORGN] OR txid11632[ORGN] OR txid10404[ORGN]) NOT phage[All Fields] NOT patent[All Fields] NOT unverified[Title] NOT vector[Title] NOT synthetic[Title] NOT chimeric[Title] NOT miRNA NOT txid28384[ORGN] NOT method[Title] NOT "partial"[Title] NOT "region"[Title] NOT "uncultured virus"[Organism] AND 00000000101[SLEN] : 00001300000[SLEN] AND ("complete genome" OR "complete segment" OR "whole genome" OR "full genome" OR "whole segment" OR "full segment") NOT txid11232[ORGN] NOT txid11233[ORGN] NOT txid82828[ORGN] NOT txid82827[ORGN] NOT txid82826[ORGN] NOT "Canine Morbillivirus" NOT "Canine Distemper" ) | 12 April 2019 |

| **Virus** | **Genotyping Region** | **Number of Sequences** | **GenBank Accession Numbers** | **Reference** |
| --- | --- | --- | --- | --- |
| BVDV | *N^pro^* | 20 | EU180034, AY182155, AY182162, AF144463, AY735490, AF287286, AF287283, AF287285, AF287279, U80902, AY894998, EU163964, KC207075, AB359929, AB359931, GU120259, KC695812, KF154777, BVU18059, AF144469 | 26 |
| CDV | *H* | 13 | LC007976, JN836734, AY465925, MH496776, EU098102, KF835413, DQ889188, FJ392651, KC916716, FJ868174, FJ461700, AY964112, HQ403645 | 27 |
| EHDV | *VP2* | 9 | JX965387, AM744988, KU140736, AM745018, AM745028, KF570134, KU523923, LC202948, AM745058 | 28 |
| IBV | *S1* | 32 | M95169, GU393336, L14069, L18988, U29522, U29519, AY606320, JQ964061, M99482, AF151954, JX182775, X52084, EU914938, X87238, FJ807932, KJ941019, AF419315, AY296744, KC577395, AF349621, DQ064806, KC577382, AF093796, KF757447, EU925393, FN182243, GU301925, M21971, U29450, U77298, DQ059618, GQ265948 | 29 |
| IAV | *HA* | 18 | CY258500, MG280408, KY284550, CY195945, KT002528, KY131360, KY013873, MG280398, MG280584, CY241348, CY206244, MG280394, CY195647, KX024583, MF145850, CY195655, CY103892, KR077932 | 30 |
|  | *NA* | 11 | KP137718, KX830941, MG280423, CY195899, KY828820, KJ568419, KY551357, KY130562, MG280527, CY103878, CY125947 |  |
| PRRSV | *ORF5* | 18 | EF031042, AY743932, M96262, EU071229, DQ324690, EU071235, U87392, EU759139, AF184212, AF494042, AF176466, AF121131, AB175721, EU757121, DQ176019, KU666088, KR154419, JQ656289 | 31 |

**Supplementary Table 2.** Information for sequences used to create custom Centrifuge indices for lineage typing viruses sequenced in this study.

BVDV= bovine viral diarrhea virus; CDV = canine distemper virus; EHDV = epizootic hemorrhagic disease virus; *H* = hemagglutinin; *HA* = hemagglutinin; IAV = influence A virus; IBV = infectious bronchitis virus; *NA* = neuraminidase; *N^pro^* = N-terminal protease; *ORF* = open reading frame; PRRSV = porcine reproductive and respiratory syndrome virus; *S1* = spike 1.
